# Impeding DNA Break Repair Enables Oocyte Quality Control

**DOI:** 10.1101/277913

**Authors:** Huanyu Qiao, H.B.D. Prasada Rao, Yan Yun, Sumit Sandhu, Jared H. Fong, Manali Sapre, Michael Nguyen, Addy Tham, Benjamin W. Van, Tiffany Y.H. Chng, Amy Lee, Neil Hunter

**Affiliations:** Howard Hughes Medical Institute, University of California, Davis, Davis, California, USA.; Department of Microbiology & Molecular Genetics, University of California, Davis, Davis, California, USA.; Department of Molecular & Cellular Biology, University of California, Davis, Davis, California, USA.; Department of Cell Biology & Human Anatomy, University of California, Davis, Davis, California, USA.

## Abstract

Oocyte quality control culls eggs with defects in meiosis. In mouse, oocyte death is triggered by defects in chromosome synapsis and recombination, which involve repair of programmed DNA double-strand breaks (DSBs) between homologous chromosomes. We show that RNF212, a SUMO ligase required for crossing over, also mediates oocyte quality control. Both physiological apoptosis and wholesale oocytes elimination in meiotic mutants require RNF212. RNF212 sensitizes cells to DSB-induced apoptosis within a narrow window when chromosomes desynapse during the transition into quiescence. Analysis of DNA damage during this transition implies that RNF212 impedes DSB repair. Consistently, RNF212 is required for HORMAD1, a negative regulator of inter-sister recombination, to associate with desynapsing chromosomes. We infer that oocytes impede repair of residual DSBs to retain a “memory” of meiotic defects that enables quality control processes. These results define the logic of oocyte quality control and suggest RNF212 variants may influence transmission of defective genomes.

## INTRODUCTION

Oocyte quality and number are important determinants of reproductive success (Kerr et al., 2013). These attributes are influenced by the selective elimination of oocytes that experience problems during the early stages of meiosis (Bolcun-Filas et al., 2014; Cloutier et al., 2015; Di Giacomo et al., 2005; Kerr et al., 2013; Kogo et al., 2012a; Malki et al., 2014; Rinaldi et al., 2017; Wojtasz et al., 2012). In mouse, epigenetic reprogramming during meiosis derepresses LINE-1 transposons triggering perinatal loss of around two thirds of all fetal oocytes in a conserved process called fetal oocyte attrition (Malki et al., 2014). Defects in the chromosomal events of meiotic prophase also trigger oocyte loss, typically at a later stage than fetal oocyte attrition, as oocytes transition into quiescence before developing into primordial follicles to establish the ovarian reserve (Di Giacomo et al., 2005). This second wave of oocyte death is mediated by interrelated pathways that signal defective interactions between pairs of homologous chromosome, i.e. defects in DSB repair and/or homolog synapsis (Bolcun-Filas et al., 2014; Cloutier et al., 2015; Di Giacomo et al., 2005; Kogo et al., 2012a; Rinaldi et al., 2017; Wojtasz et al., 2012). Canonical DNA-damage response factors signal unrepaired DSBs via the ATR and CHK2 kinases to trigger p53/p63-mediated oocyte apoptosis. Defective synapsis can also trigger oocyte death by a distinct pathway that mediates transcriptional silencing in a process termed meiotic silencing of unpaired chromatin (MSUC). Together, the perinatal and pre-follicle oocyte elimination pathways balance the quality and size of the ovarian follicle reserve to maximize reproductive success. In this study, we address why and how defective inter-homolog interactions trigger oocyte apoptosis after chromosomes desynapse during the transition into quiescence. We implicate a new factor, RNF212, in the pre-follicle oocyte apoptosis pathway and show that it functions in a counterintuitive process that helps oocytes gauge whether meiotic inter-homolog interactions were defective.

## RESULTS

### *Rnf212* Mutants Have Enlarged Oocyte Reserves

RNF212 is a RING-family E3-ligase that regulates the progression of meiotic recombination via the SUMO modification and ubiquitin-proteasome systems, and is essential for crossing over (Qiao et al., 2014; Rao et al., 2017; Reynolds et al., 2013). *Rnf212* mutation in mice causes sterility in both sexes. Although early events of meiotic prophase occur efficiently in *Rnf212*^−/−^mutants, including synapsis and DSB repair, crossing-over fails leaving homologs unconnected at metaphase I. In males, the resulting univalents are a potent trigger of apoptosis resulting in the complete absence of spermatids (Reynolds et al., 2013). Elimination of gametes was not seen in *Rnf212*^−/−^ mutant females (**Figure 1, A–E**). On the contrary, large numbers of oocytes were present in *Rnf212*^−/−^ mutant ovaries. This sexually dimorphic response to *Rnf212*^−/−^ mutation presumably reflects the fact that postnatal oocytes uniformly arrest in the dictyate stage, which follows crossing over and desynapsis of homologous chromosomes, but precedes meiosis **I**.Thus, the absence of crossovers in *Rnf212*^−/−^ mutant oocytes goes undetected until they resume meiosis in the preovulatory stage of folliculogenesis, as seen for other mutants defective for late steps of crossing over (Kan et al., 2008). Consistent with this interpretation, *in vitro* maturation experiments showed that oocytes from *Rnf212*^−/−^ mutants resume meiosis efficiently, but fail to successfully execute meiosis **I** (**Figure S1**). Thus, the absence of early defects in DSB repair and homolog synapsis allows establishment of non-fertile follicle reserves in *Rnf212*^−/−^ mutants.

**Figure 1:**
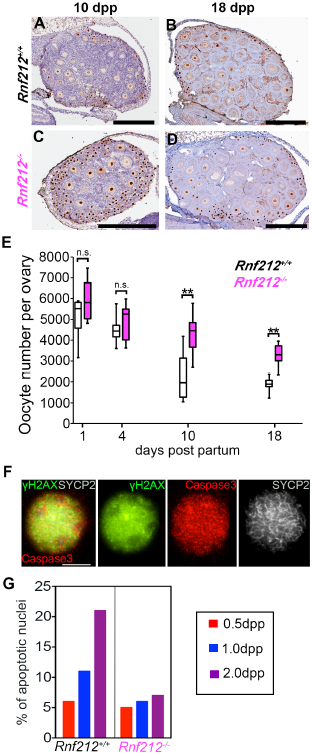
**RNF212 promotes postnatal oocyte apoptosis**. (**A**, **B**) Representative wild type and (**C**, **D**) *Rnf212* mutant ovary sections from 10 and 18 dpp females immunostained for p63 with hematoxylin counterstaining. (**E**) Oocyte counts in wild type and *Rnf212*^−/−^ mutant ovaries. Box plots show 25th and 75th percentiles (box), median (the line in the center of the box), and minimum and maximum values (whiskers). \*\**P* ≤ 0.01, two-tailed Mann-Whitney test. (**F**) Representative wild-type oocyte nucleus at 2 dpp immunostained for γH2AX (green), caspase 3 (red) and chromosome axis marker SYCP2 (grey). (**G**) Quantification of apoptotic nuclei at successive days postpartum. Scale bar in F. is 10μm.

Quantification of oocyte numbers revealed that reserves were significantly larger in *Rnf212*^−/−^ mutants than in their wild-type counterparts. Ovaries dissected from 1, 4, 10 and 18 day old animals (day post-partum, dpp) were fixed, sectioned and immunostained for oocyte-specific markers (MVH, SYCP3 and/or p63)(**Figure 1, A–D,** and **Figure S2**). At 1 and 4 days post partum (dpp), oocyte numbers in *Rnf212*^−/−^ mutant ovaries appeared slightly increased relative to their wild-type counterparts, but differences were not statistically significant (**Figure 1E**). However, the ovaries of 10 and 18 dpp *Rnf212*^−/−^ mutants contained 45-57% more oocytes than wild-type ovaries.

### RNF212 Promotes Physiological and Pathological Oocyte Apoptosis

One explanation for the larger oocyte pools of *Rnf212*^−/−^ mutants is that RNF212 promotes pre-follicle oocyte apoptosis. To address this possibility, we immunostained early postnatal oocytes from 0.5, 1.0 and 2.0 dpp animals for the DNA-damage marker, phosphorylated histone H2AFX (γH2AX), and the apoptosis execution factor, Caspase 3 (**Figure 1F** and **1G**). A subset of oocyte nuclei was identified with pan-nuclear staining for both γH2AX and Caspase 3, which is diagnostic of apoptotic nuclei (Harada et al., 2014) (**Figure 1F**). In wild-type animals, the fraction of apoptotic oocytes increased ~4-fold between 0.5 and 2.0 dpp (from 6 to 21%) indicating elevated apoptosis as oocytes transitioned into dictyate (**Figure 1G**). This increase was not seen in the *Rnf212*^−/−^ mutant, for which levels of apoptotic nuclei remained low and relatively constant, at ~5% of total oocytes, between birth and 2 dpp. We conclude that RNF212 facilitates the apoptosis of a subset of early postnatal oocytes.

To test the idea that RNF212 promotes apoptosis of oocytes that have experienced errors in meiotic prophase I, we asked whether *Rnf212* mutation could rescue the wholesale elimination of oocytes caused by *Spo11* and *Msh4* mutations (**Figure 2**). *Spo11*^−/−^ mutants fail to initiate meiotic recombination resulting in severely defective homolog synapsis and almost complete elimination of the oocyte pool (**Figure 2, A–C**) (Baudat et al., 2000). Consistent with previous analysis (Di Giacomo et al., 2005), only a very small number of oocytes survived in *Spo11*^−/−^ mutants at 18 dpp, less than 7.5% of wild type (**Figure 2A** and **1C**). However, in *Spo11*^−/−^ *Rnf212*^−/−^ double mutants, large numbers of oocytes were present, averaging 40% of the oocyte numbers seen in *Rnf212*^−/−^ single mutants (**Figure 2B, 2C** and **2G**). More complete restoration of oocyte pools by *Rnf212* mutation was seen in the *Msh4*^−/−^ mutant background. Absence of MSH4 causes severe defects in both DSB repair and homolog synapsis resulting in the complete elimination of oocytes within 4 days of birth (Kneitz et al., 2000)(**Figure 2 D** and **2F**).

**Figure 2:**
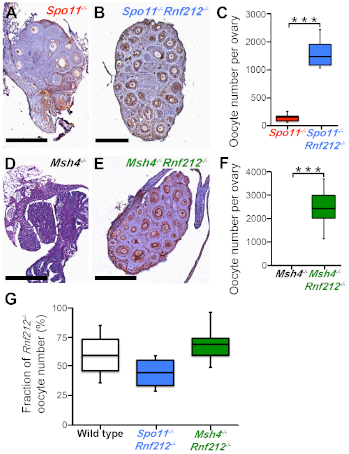
RNF212 is required for wholesale oocyte apoptosis in response to defects in early meiotic prophase. (**A**) Representative *Spo1*^−/−^and (**B**) *Spo11*^−/−^*Rnf212*^−/−^18 dpp ovary sections immunostained for p63 with hematoxylin counterstaining. (**C**) Oocyte counts in *Spo11*^−/−^and *Spo11*^−/−^*Rnf212*^−/−^ovaries at 18 dpp. (**D**) Representative *Msh4*^−/−^and (**E**) *Rnf212*^−/−^*Msh4*^−/−^18 dpp ovary sections. The dashed line highlights the degenerate ovary in the *Msh4*^−/−^mutant. (**F**) Oocyte counts at 18 dpp. (**G**) Oocyte counts at 18 dpp in wild type, *Spo11*^−/−^*Rnf212*^−/−^and *Msh4*^−/−^*Rnf212*^−/−^expressed as a percentage of *Rnf212*^−/−^oocyte numbers. \*\*\**P* ≤ 0.001, two-tailed Mann-Whitney test. Scale bars for ovary sections are 300 μm.

Strikingly, the ovaries of *Msh4*^−/−^ *Rnf212*^−/−^ double mutants contained high numbers of oocytes, averaging 75% of those in the *Rnf212*^−/−^ single mutant (**Figure 2E, 2F** and **2G**). Moreover, small numbers of oocytes were able to escape apoptosis even when *Rnf212* was heterozygous in the *Msh4*^−/−^ background (**Figure S3**). Thus, the complete elimination of oocytes in *Msh4*^−/−^ mutants is sensitive to the level of RNF212.

### RNF212 Modulates DNA Damage Levels in Early Postnatal Oocytes

The data above indicate that RNF212 promotes apoptosis of oocytes that have experienced defects in meiotic prophase I. RNF212 could influence the primary signaling of defects in recombination and synapsis, for example by altering DSB repair. Alternatively, RNF212 could be a component of downstream signaling pathways that ultimately trigger apoptosis. To distinguish these possibilities, we monitored levels of DNA damage in early postnatal oocyte nuclei via γH2AX immunostaining (**Figure 3A** and **3B**). During meiosis, γH2AX decorates the chromatin around DSBs and accumulates at regions of chromosomes that fail to undergo synapsis (Cloutier et al., 2015; Mahadevaiah et al., 2001). We reasoned that if RNF212 functions solely in downstream steps to signal damage or trigger apoptosis, presumptively defective oocytes with high levels of γH2AX should be present at highly elevated levels in the ovaries of *Rnf212*^−/−^ mutants. Oocytes from 1 dpp mice were found to be in one of three stages: late pachytene, diplotene or early dictyate (**Figure 3A** and **Figure S4**). In late pachytene nuclei, an average of 40 ± 15 γH2AX chromatin “flares” were detected, decreasing to 21 ± 11 in diplotene, and just 6 ± 3 as oocytes transitioned into dictyate (**Figure 3A** and **3B**). Contrary to expectations, numbers of γH2AX flares were ~2-fold lower at all stages in the early postnatal oocytes of *Rnf212*^−/−^ mutants implying lower DSB levels. *Rnf212*^−/−^ oocytes entering dictyate contained an average of only 2.7 ± 1.4 γH2AX flares, compared to 6 ± 3 in wild-type cells; moreover, 20% of nuclei had no staining at all, compared to just 6% of wild-type oocytes (**Figure 3A** and **3B**).Qualitatively, the γH2AX signals detected in *Rnf212*^−/−^ oocytes were generally fainter and more compact than those seen in wild type.

**Figure 3:**
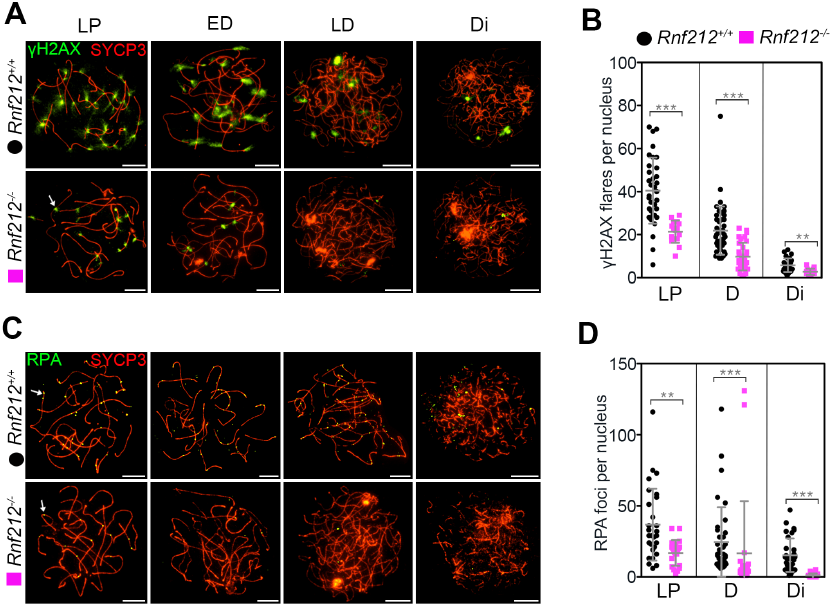
RNF212 impedes DNA-damage repair in early-postnatal oocytes. Representative 1 dpp oocyte nuclei at the indicated stages immunostained for chromosome axis marker SYCP3 (red) and γH2AX (green). Arrow indicates an individual γH2AX flare. (**B**) Quantification of γH2AX flares in oocyte nuclei at successive prophase substages. (**C**) Representative 1 dpp oocyte nuclei at the indicated stages immunostained for chromosome axis marker SYCP3 (red) and RPA (green). Arrows highlight individual RPA foci. (**D**) Quantification of RPA foci in 1 dpp oocyte nuclei from the indicated strains immunostained for chromosome axis marker SYCP3 (red) and RPA (green).Arrows highlight individual RPA foci. (**E**) Quantification of RPA foci in 1 dpp oocytes from the indicated strains. \*\**P* ≤ 0.01, \*\*\**P* ≤ 0.001, two-tailed Mann-Whitney test. Error bars show means ± SD. Scale bars, 10μm.

Given that γH2AX can mark both DSBs and non-DSB lesions during meiosis (Cloutier et al., 2015), we also immunostained oocytes from 1 dpp animals for Replication Protein A (RPA), which tightly binds regions of single-stranded DNA created during replication, repair and recombination (Ribeiro et al., 2016)(**Figure 3C** and **3D**). RPA-staining foci were detected in similar numbers to γH2AX flares, suggesting that the vast majority of γH2AX signals in wild-type and *Rnf212*^−/−^ oocytes mark sites of DNA lesions, very likely DSBs. Moreover, like γH2AX flares, numbers of RPA foci were reduced ~2-fold in *Rnf212*^−/−^ mutant oocytes. Together, these data imply that RNF212 modulates the generation or processing of DNA damage that leads to oocyte apoptosis.

One possibility is that RNF212 somehow impedes DSB repair in early postnatal oocytes. This idea was further explored by quantifying DNA-damage markers in early postnatal oocytes of the meiotic mutants, *Spo11*^−/−^ and *Msh4*^−/−^, and the corresponding double mutants with *Rnf212*^−/−^ (**Figure 4**). The defective homolog synapsis that occurs in *Spo11*^−/−^ and *Msh4*^−/−^ mutants makes it hard to distinguish oocytes in late pachytene from those in diplotene. Therefore, we could assign only two categories of oocyte nuclei in these mutants: “early” nuclei had bright, contiguous SYCP3-staining axes and were inferred to be in pachytene and diplotene-like stages; and “late” nuclei had fainter, fragmented SYCP3 axes and were inferred to be either early dictyate or early apoptotic oocytes (**Figure 4A**). In early-stage *Spo11*^−/−^ oocytes from 1 dpp animals, γH2AX staining was present as numerous large flares, encompassing long portions of individual chromosome axes indicative of the extensive transcriptional silencing that occurs in response to asynapsis called meiotic silencing of unpaired chromatin (MSUC)(Baarends et al.,2005; Cloutier et al., 2015; Kouznetsova et al., 2009; Turner et al., 2005). 88% of *Spo11*^−/−^ oocytes showed this pattern, while 12% had pan-nuclear staining (**Figure 4A, 4B** and **4C,** and **Figure S4**). Much more extensive γH2AX staining was seen in late-stage *Spo11*^−/−^ mutant oocytes; more than 91% showed pan-nuclear staining, while the remainder had a pattern of extended γH2AX flares. Mutation of *Rnf212* in the *Spo11*^−/−^ background greatly reduced the extent of γH2AX staining. In late-stage *Rnf212*^−/−^ *Spo11*^−/−^ oocytes, the fraction of nuclei with pan-nuclear staining was reduced by half (45% versus 91% in the *Spo11*^−/−^ single mutant) with a corresponding increase in nuclei with γH2AX flares. Moreover, for both early and late-stage *Rnf212*^−/−^ *Spo11*^−/−^ oocytes containing γH2AX flares, the total staining areas were greatly reduced, by 5-fold, relative to those of nuclei from the *Spo11*^−/−^ single mutant (**Figure 4E** and **Figure S4**). *Rnf212* mutation also diminished γ H2AX signals in *Msh4*^−/−^ mutant oocytes, but the effects were more extreme (**Figure 4A, 4B** and **4C,** and **Figure S4**). 85% of early-stage and 91% of late-stage oocytes from the *Msh4*^−/−^ mutant showed pan-nuclear γ H2AX staining. Oppositely, ~80% of oocytes from *Rnf212*^−/−^ *Msh4*^−/−^ double mutants had the γH2AX flare-staining pattern, with greatly reduced (30-fold) staining areas compared to those of *Msh4*^−/−^ single mutant nuclei. Thus, *Rnf212* mutation diminishes the very high levels of γ H2AX that occur in early postnatal oocytes of both *Spo11*^−/−^ and *Msh4*^−/−^ mutants.

**Figure 4:**
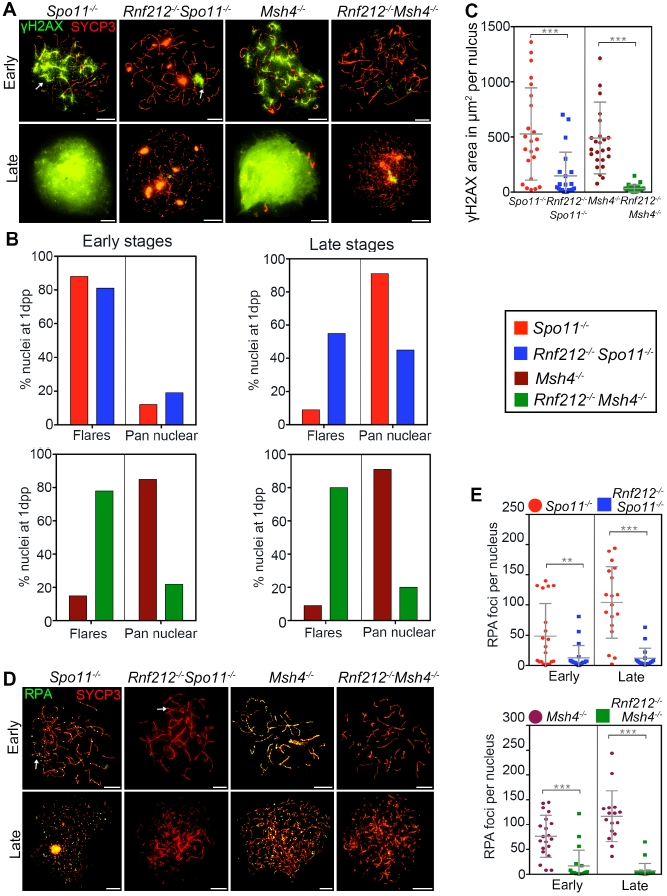
RNF212 impedes DNA-damage repair in *Spo11* and *Msh4* mutants. **(A)** Representative 1 dpp oocyte nuclei from the indicated strains immunostained for chromosome axis marker SYCP3 (red) and γH2AX (green). (**B**) Quantification of γH2AX flares and pan nuclear γH2AX staining in 1 dpp oocytes. (**C**) Quantification of γH2AX immunostaining area in 1 dpp oocytes. (**D**) Representative 1 dpp oocyte nuclei from the indicated strains immunostained for chromosome axis marker SYCP3 (red) and RPA (green). Arrows highlight individual RPA foci. (**E**) Quantification of RPA foci in 1 dpp oocytes from the indicated strains. \*\**P* ≤ 0.01, \*\*\**P* ≤ 0.001, two-tailed Mann-Whitney test. Error bars show means ± SD. Scale bars, 10μm.

Supporting the interpretation that *Rnf212* mutation is reducing the level of DNA damage present in early postnatal oocytes of *Spo11*^−/−^ and *Msh4*^−/−^ mutants, RPA foci showed changes similar to those seen for γH2AX (**Figure 4D** and **4E**). In *Spo11*^−/−^ and *Msh4*^−/−^ single mutants, early-stage oocytes contained an average of 50 ± 53 and 77 ± 42 RPA foci, respectively. The presence of high levels of Spo11-independent DSBs in *Spo11*^−/−^ mutant oocytes was previously documented and shown to make a significant contribution to oocyte apoptosis in this asynaptic background via the DNA damage response (Carofiglio et al., 2013; Carofiglio et al., 2018; Rinaldi et al., 2017). Although the origin of these breaks is currently unknown, a likely source is LINE-1 activity (Malki et al., 2014). In contrast to wild-type animals, in which RPA focus numbers decreased as oocytes transitioned into dictyate (**Figure 3C** and **3D**), foci further increased in late-stage *Spo11*^−/−^ and *Msh4*^−/−^ mutant oocytes (**Figure 4D** and **4E**), mirroring the transition to pan-nuclear γH2AX staining (**Figure 4A** and **4B**). *Rnf212* mutation decreased RPA focus numbers by 3 to 4 fold in both *Spo11*^−/−^ and *Msh4*^−/−^ backgrounds. Thus, consistent with the idea that RNF212 impedes DSB repair in early postnatal oocytes, the presence of RNF212 greatly enhances DNA damage levels in of meiotic mutants defective for homolog synapsis and DSB repair.

### RNF212 is Required for HORMAD1 to Associate With Desynapsing Chromosomes as Oocytes Transition into Quiescence

The ability of *Rnf212* mutation to suppress the oocyte death of recombination and synapsis mutants is shared by mutation of the *Hormad1* gene that encodes a meiosis-specific HORMA (Hop1, Rev7 and Mad2)-domain protein with central roles in regulating the events of meiotic prophase I (Daniel et al., 2011; Kogo et al., 2012b; Shin et al., 2013). HORMAD1 and its orthologs have been implicated in meiotic DSB formation, homolog pairing and synapsis, checkpoint signaling, transcriptional silencing (MSUC), and biasing meiotic recombination to occur between homologs by impeding inter-sister DSB repair (Carballo et al., 2008; Rinaldi et al., 2017; Royo et al., 2013; Shin et al., 2010; Stanzione et al., 2016; Vader and Musacchio, 2014). HORMAD1 is initially associated with unsynapsed chromosome axes during early prophase I, but then becomes depleted as homologs synapse, before re-associating with desynapsing axes during diplonema (Fukuda et al.; Niu et al., 2005; Wojtasz et al., 2009)(**Figure 5A**).

**Figure 5:**
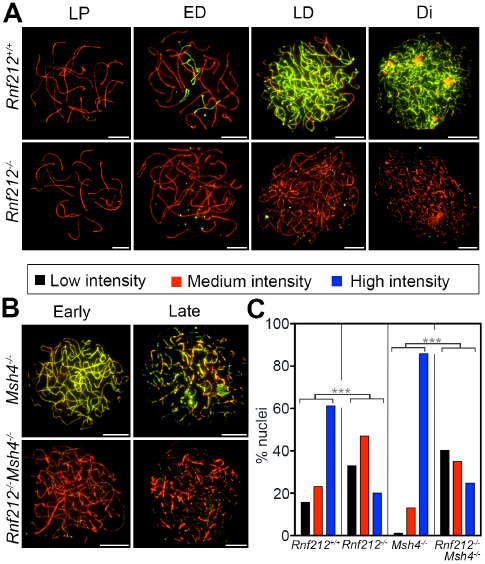
RNF212 promotes association of HORMAD1 with desynapsed homolog axes. (**A**, **B**) **Representative** 1 dpp oocyte nuclei from the indicated strains immunostained for SYCP3 (red) and HORMAD1 (green). LP, late pachynema; ED, early diplonema; LD, late diplonema; Di, early dictyotene. (**C**) Quantification of HORMAD1 signal intensity ratios in the indicated strains. For each nucleus, total signal for HORMAD1 was divided by total signal for SYCP3: ratio of ≤0.2, low intensity; 0.21-0.4, medium intensity; ≥0.41, high intensity. \*\*\**P* ≤ 0.001, G-test. Scale bars, 10μm.

The role of HORMAD1 upon its reassociation with desynapsed diplotene chromosomes is unknown, but its timing and location suggest the potential to influence the repair of residual DSBs between sister chromatids. Therefore, we asked whether *Rnf212* mutation affects the re association of HORMAD1 with diplotene and early dictyate chromosomes (**Figure 5A**). Immunostaining of oocyte chromosome spreads from 1 dpp wild-type animals revealed strong association of HORMAD1 with desynapsed homolog axes in early diplotene nuclei (**Figure 5A**, top row). In nuclei where desynapsis was more advanced, extensive association of HORMAD1 was seen along the lengths of all desynapsed axes. This extensive HORMAD1 staining pattern persisted into early dictyate, even in nuclei where the SYCP3-staining axes had become faint and highly fragmented. In *Rnf212*^−/−^ mutant oocytes, association of HORMAD1 with diplotene and dictyate chromosomes was greatly diminished (**Figure 5A**, bottom row). To quantify this defect, we binned nuclei into three HORMAD1 staining-classes based on their relative signal intensities (**Figure 5C**). In wild-type nuclei from 1 dpp animals, 61, 23 and 16% of nuclei had high, medium and low intensity HORMAD1 staining, respectively; compared to 20, 47 and 33% of *Rnf212*^−/−^ mutant nuclei (*P*<0.001, G-test). We also examined HORMAD1 dynamics in 1 dpp *Msh4*^−/−^ mutants, in which all oocytes are destined for apoptosis, and in *Rnf212*^−/−^ just 24% of nuclei from *Rnf212*^−/−^*Msh4*^−/−^double mutants where most oocytes survive (**Figure 5B** and **5C**). High-intensity HORMAD1 staining was observed in 85% of *Msh4*^−/−^ oocyte nuclei (both early and late stage nuclei, defined by the criteria described above). Again, chromosomal association of HORMAD1 was greatly diminished by *Rnf212*^−/−^ mutation; just 24% of nuclei from *Rnf212*^−/−^*Msh4*^−/−^double mutants showed high intensity HORMAD1 staining (*P*<0.001, G-test). Thus, RNF212 promotes or stabilizes the association of HORMAD1 along desynapsed homolog axes both in wild-type oocytes and in mutants such as *Msh4*^−/−^that experience severe defects in synapsis and recombination.

### RNF212 Enhances DSB-Induced Apoptosis as Oocytes Transition into Quiescence

The data above suggest that the transition into quiescence, from the diplotene to dictyate stages, marks a critical oocyte quality-control juncture where the success of meiotic prophase is determined by assessing residual levels of DNA damage. To test this interpretation we asked if exogenous DSBs, inflicted specifically during the early postnatal period, could trigger oocyte apoptosis and whether RNF212 enhanced this process (**Figure 6**). Animals were exposed to γ-irradiation at different times during the two-day window after birth when oocytes transition into dictyate. Previous studies showed that exposure of 5 dpp mice to 0.45 Gy, estimated to induce ~10 DSBs per nucleus, efficiently triggered apoptosis (Rinaldi et al., 2017; Suh et al., 2006). Therefore, we initially exposed wild-type and *Rnf212*^−/−^littermates, at 0.5, 1 or 2 dpp, to 0.45 Gy. Oocyte numbers from irradiated mice were then quantified at 10 dpp and compared to unirradiated controls (**Figure 5A–D**). At 0.5 dpp, oocytes were largely resistant to radiation-induced apoptosis. Oocyte numbers in both wild-type and *Rnf212*^−/−^ animals were similar to those of unirradiated wild-type controls (*P>* 0.58; **Figure 6D**), but still smaller than those of unirradiated *Rnf212*^−/−^animals. In contrast, oocytes at 1 and 2 dpp were highly sensitive to radiation-induced apoptosis. Survival of wild-type oocytes averaged only 16% and 7% of unirradiated controls following exposure to 0.45 Gy at 1 and 2 dpp, respectively (**Figure 6D**). Survival of *Rnf212*^−/−^ oocytes irradiated at 1 and 2 dpp was slightly higher than wild type, averaging respectively 21% and 10% of the *Rnf212*^−/−^mutant unirradiated controls (or 39% and 20% of wild-type unirradiated controls). Thus, at 0.5 dpp, both wild-type and *Rnf212*^−/−^oocytes are largely resistant to a dose of irradiation that becomes a potent trigger of apoptosis at 1 and 2 dpp, as oocytes transition into dictyate.

**Figure 6:**
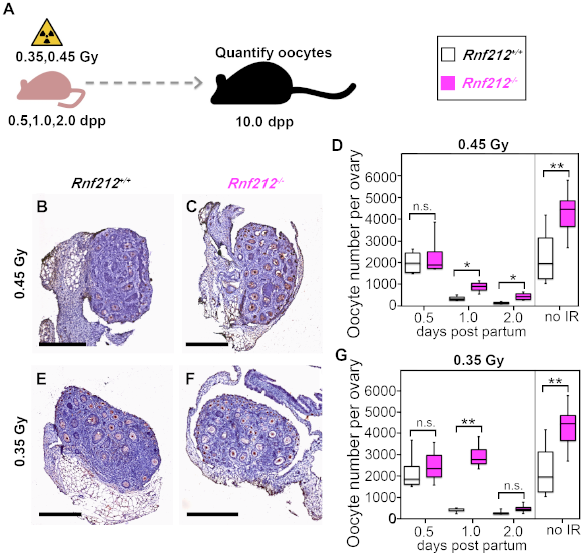
RNF212 sensitizes early-postnatal oocytes to radiation-induced apoptosis. (**A**) Experimental regimen for irradiation experiments. (**B**, **C** and **E**, **F**) 10 dpp ovary sections from *Rnf212*^−/−^and *Rnf212*^−/−^females following irradiation with 0.45 or 0.35 Gy at 1 dpp. (**D** and **G**) Oocyte counts following irradiation with 0.45 or 0.35 Gy, respectively. (**G**) Quantification of oocyte numbers following irradiation with 0.45 Gy. Box plots show 25th and 75th percentiles (box), median (the line in the center of the box), and minimum and maximum values (whiskers).\**P* ≤ 0.05, \*\**P* ≤ 0.01, \*\*\**P* ≤ 0.001, two-tailed Mann-Whitney test. Scale bars, 300 μm.

Suh et al. (Suh et al., 2006) demonstrated a narrow dose-response relationship between γ-irradiation and oocyte apoptosis. This fact, together with the suggestion that *Rnf212*^−/−^mutant oocytes are slightly more resistant to 0.45 Gy at 1 and 2 dpp (**Figure 6D**), motivated us to examine the effects of a lower dose of irradiation (**Figure 6E–G**). With a dose of 0.35 Gy (equivalent to ~7 DSBs per nucleus), both wild-type and *Rnf212*^−/−^oocytes were again largely resistant to apoptosis when exposed at 0.5 dpp, but highly sensitive when exposed at 2 dpp, similar to the responses observed with 0.45 Gy. However, a striking difference in sensitivity was observed when mice were exposed at 1 dpp. While wild-type oocytes remained highly sensitive, averaging only 13% survival (relative to unirradiated wild type), *Rnf212*^−/−^mutant oocytes were largely resistant with 68% survival relative to unirradiated *Rnf212*^−/−^controls (**Figure 6G**). We conclude that RNF212 enhances DSB-induced apoptosis during a narrow window as oocytes transition from diplotene to dictyate.

## DISCUSSION

Together, our data delineate a major quality control point at the diplotene-to-dictyate transition during which oocytes actively impede DSB repair in order to gauge whether the events of early meiotic prophase occurred successfully (**Figure 7**). At this stage, an unrepaired DSB is indicative of a failed inter-homolog interaction, or possibly a lesion resulting from LINE-1 activity (Malki et al., 2014). During diplotene, homologs are no longer closely juxtaposed (except very locally at crossover sites) and inter-homolog recombination factors are no longer present such that accurate repair of residual DSBs can only be achieved via recombination with the sister chromatid. By blocking inter-sister recombination, RNF212-dependent re-loading of HORMAD1 onto desynapsed homolog axes effectively acts to preserve a “memory” of meiotic prophase errors. Reinstallation of HORMAD1 could similarly impede repair of LINE-1 induced damage, and also reinforce transcriptional silencing via MSUC, which requires HORMAD proteins (Royo et al., 2013; Wojtasz et al., 2012). Consequently, a robust signaling response from any residual lesions will be maintained during the diplotene-to-dictyate transition enabling a meiotic quality-control decision to be made (**Figure 7**). Signaling above a critical threshold efficiently triggers apoptosis preventing oocytes that have experienced significant recombination and synapsis errors, or high levels of LINE-1 activity, from contributing to the ovarian reserve. Consistent with the study of Rinaldi et al. (2017), we estimate that this threshold is equivalent to ~10 DSBs per nucleus (**Figure 6**). Importantly, our data indicate that pre-follicle apoptosis is a significant physiological quality-control pathway that removes around half of the oocytes that remain after LINE-1 induced fetal oocyte attrition (Malki et al., 2014). Together, we can estimate that fetal and pre-follicle attrition cull over 80% of all oocytes in mouse (Malki et al., 2014)(this study), mirroring the dramatic reduction in oocyte numbers seen in human females between ~20 weeks gestation and birth (Findlay et al., 2015), presumably via equivalent quality control mechanisms.

**Figure 7:**
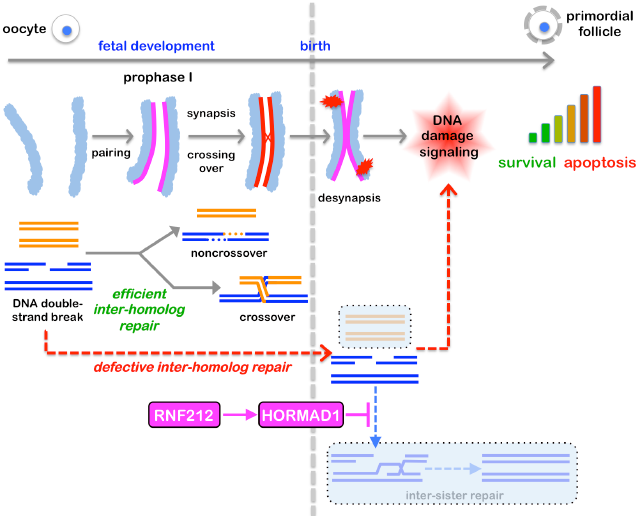
Logic of pre-follicle oocyte quality control. Timeline of chromosomal and DNA events during oocyte meiotic prophase. Magenta lines indicate homolog axes associated with HORMAD1. Red “explosions” represent unrepaired DSBs. Transparent boxes indicate repair pathways that are unavailable following desynapsis in diplotene. Inter-sister repair is actively impeded by the RNF212-dependent reassociation of HORMAD1 with desynapsed homolog axes such that any residual DSBs robustly signal to the checkpoint machinery. The integrated level of damage signaling from all residual DSBs dictates whether an individual oocyte undergoes apoptosis or survives to form a primordial follicle and become part of the ovarian reserve.

Analysis of recombination intermediates in budding yeast has provided direct evidence that HOP1^HOMARD^ negatively regulates inter-sister recombination as part of a phospho-kinase signal-transduction pathway that responds to DSBs (Humphryes and Hochwagen, 2014; Lao and Hunter, 2010). Following DSB-dependent phosphorylation by PI3K-like sensor kinases, Mec1^ATR^ and Tel1^ATM^,Hop1^HOMARD^ acts as a mediator for activation of the effector kinase, Mek1, which directly targets components of the recombination and cell-cycle machinery (Callender et al., 2016; Carballo et al., 2008; Prugar et al., 2017). Two lines of evidence in mouse indicate that mammalian HORMADs also negatively regulate inter-sister recombination; first, repair of irradiation-induced DSBs is accelerated when HORMADs are absent; and second,DSB levels are diminished in asynaptic *Spo1*^−/−^mutant meiocytes in which inter-sister recombination is likely the only repair option (Carofiglio et al., 2018; Rinaldi et al., 2017; Shin et al., 2013).

HORMAD1 initially loads onto leptotene chromosomes independently of RNF212 (data not shown). After homolog pairing has been achieved, we propose that nascent interhomolog interactions are “locked in” by synaptonemal complexes and associated meiosis-specific recombination factors (ZMMs)(Hunter, 2015; Lynn et al., 2007). Consequently, the role of HORMAD proteins in preventing intersister recombination is no longer necessary. Indeed, dissociation of HORMADs as synapsis ensues may enable inter-sister repair of DSBs that either failed to engage a homolog template, or that engaged the interhomolog DNA strand exchange to facilitate pairing, but did not complete repair. Estimates of the numbers of unrepaired DSB numbers in the *Trip13* mutant, which fails to remove HORMADs from homolog axes upon synapsis, implies that up to a quarter of DSBs may utilize inter-sister recombination for their repair (Rinaldi et al., 2017). Our observation that late pachytene cells are relatively insensitive to DSB-induced apoptosis (**Figure 6**) could be explained, at least in part, by a larger capacity to repair DSBs via inter-sister recombination following the dissociation of HORMAD proteins. As such, many more DSBs would be required to reach the signaling threshold required to induce apoptosis at this stage. In addition, the signaling threshold required to induce apoptosis during pachytene may be increased in order to prevent aberrant culling of oocytes that are still completing recombination (Kim and Suh, 2014). The window in which RNF212 enhances DSB-induced apoptosis also likely reflects the timing of expression of the transactivating p63 isoform, TAp63. As oocytes progress into dictyate, TAp63, becomes fully expressed and activatable by phosphorylation rendering primordial follicles exquisitely sensitive to DNA damage (Kim and Suh, 2014; Livera et al., 2008; Suh et al., 2006).

RNF212 is known for its essential role in meiotic crossing over (Qiao et al., 2014; Rao et al., 2017; Reynolds et al., 2013). At the cytological level, RNF212 promotes chromosome-associated SUMO conjugation during early meiotic prophase and renders the progression of recombination beyond nascent strand-exchange intermediates dependent on the action of the ubiquitin-proteasome system, mediated specifically by a second RING ligase, HEI10. RNF212 initially localizes as numerous puncta along synapsed chromosomes during zygotene and early pachytene, before accumulating specifically at future crossover sites (Reynolds et al., 2013). RNF212 is not detected along desynapsing chromosomes during diplotene, i.e. during the time it mediates the reassociation of HORMAD1. This observation suggests that RNF212 functions in the nucleoplasm to license HORMAD1 to reassociate with homolog axes upon desynapsis. Possibly, RNF212 antagonizes the action of TRIP13, an AAA+ ATPase required to dissociate HORMADs from synapsed homolog axes (Li and Schimenti, 2007; Roig et al., 2010; Wojtasz et al., 2009; Ye et al., 2017). Given that low levels of chromosome-associated HORAMD1 are still detected in *Rnf212* mutant oocytes during diplotene and dictyate (**Figure 5**), RNF212 may primarily affect the ability of HORMAD1 to oligomerize, a fundamental attribute of HORMAD proteins that is central to their function (Kim et al., 2014; Rosenberg and Corbett, 2015; Ye et al., 2017). Like yeast Hop1, HORMAD1 is important for DSB formation, thus *Hormad1* mutant phenotypes reflect both reduced DSBs and derepression of inter-sister recombination (Daniel et al., 2011; Shin et al., 2010). *Rnf212* mutation does not impact early roles of HORMAD1 in DSB formation and inter-homolog bias (Reynolds et al., 2013), but reveals a previously undefined late role as it reassociates with diplotene chromosome axes. This unanticipated role of RNF212 implicates SUMO modification in oocyte quality control and suggests that common variants of mammalian *Rnf212* genes (Johnston et al., 2016; Kadri et al., 2016; Kong et al., 2008) could have compound effects on crossing over, and the size and quality of oocyte reserves.

## ACKNOWLEDGMENTS

We thank Richard Shultz for advice and support; Vera Rinaldi, Ewilina Bolcun-Filas and John Schimenti for discussions and sharing unpublished data; Attila Toth, Christer Höög and Scott Keeney for antibodies; Jim Trimmer for ScanScope access; Shanie McCarty for assistance with irradiation experiments; and the Hunter Lab for support and discussions. H.Q. was supported by NICHD K99 Pathway to Independence Award K99HD082375. S.S. was supported by the A.P. Giannini Foundation. This work was initially supported by NIGMS grant GM084955. N.H. is an Investigator of the Howard Hughes Medical Institute.

## AUTHOR CONTRIBUTIONS

H.Q., H.B.D.P.R. and N.H. conceived the study and designed the experiments. H.Q., H.B.D.P.R., F.H.F., M.S., M.N., A.T., B.W.V., T.Y-H.C. and A.L. performed the experiments and analyzed the data. S.S. advised and contributed to histological analysis. Y.Y. performed oocyte culture experiments. H.Q., H.B.D.P.R. and N.H. wrote the manuscript with inputs and edits from all authors.

